# Customization of a *dada2*-based pipeline for fungal Internal Transcribed Spacer 1 (ITS 1) amplicon datasets

**DOI:** 10.1101/2021.07.19.452952

**Authors:** Thierry Rolling, Bing Zhai, John V. Frame, Tobias M. Hohl, Ying Taur

**Affiliations:** Infectious Disease Service, Department of Medicine, Memorial Sloan Kettering Cancer Center, New York, NY, USA; Immunology Program, Sloan Kettering Institute, Memorial Sloan Kettering Cancer Center, New York, NY, USA; Division of Infectious Diseases, First Department of Medicine, University Medical Center Hamburg-Eppendorf, Hamburg, Germany; Weill Cornell Medical College, New York, NY, USA

## Abstract

Identification and analysis of fungal communities commonly rely on internal transcribed spacer (ITS)-based amplicon sequencing. Currently, there is no gold standard to infer and classify fungal constituents, in part since methodologies have been adapted from analyses of bacterial communities. To achieve high resolution inference of fungi in clinical samples, we customized a DADA2-based pipeline using a mock community of eleven medically relevant fungi. While DADA2 allowed the discrimination of ITS1 sequences differing by a single nucleotide, quality filtering, sequencing bias, and database selection were identified as key variables determining the accuracy of sample inference. By fine-tuning quality filtering, we decreased the number of wrongly discarded sequences attributed to *Aspergillus* species, *Saccharomyces cerevisiae*, and *Candida glabrata* reads. We confirmed this effect in patient samples. By adapting a wobble nucleotide in the ITS1 forward primer region, we further increased the yield of *S. saccharomyces* and *C. glabrata* sequences. Finally, we showed that a BLAST-based algorithm based on the UNITE+INSD or the NCBI NT database achieved a higher reliability in species-level taxonomic annotation than the naïve Bayesian classifier implemented in DADA2. These steps optimized a robust fungal ITS1 sequencing pipeline that, in most instances, enables species level-assignment of community members.

## Introduction

Amplicon-based sequencing methods have allowed researchers to dissect the composition of the bacterial microbiota in a broad range of environmental and biological samples and have widened our knowledge of host-microbe interactions in health and disease (1). More recently the target of microbiota research has expanded beyond the bacterial kingdom to encompass fungi, archaea, and viruses. The recognized target for fungal taxonomic profiling is the internal transcribed spacer region of the rDNA (ITS)(2). Due to the sequencing length limitations, only one of the two subregions ITS1 or ITS2 is commonly used. We have previously shown the potential of an ITS1-based approach to identify the intestinal origin of *Candida* bloodstream infections (3). Here, we show how to customize this ITS1-based platform to more accurately reflect species representation.

Currently, most of the amplicon-based microbiota profiling methods rely on sequencing using an Illumina platform. Starting with the raw Illumina sequences, a pipeline for amplicon analysis includes multiple steps: (optional) demultiplexing, primer removal, quality filtering, denoising or OTU picking, and taxonomic annotation. The components of these pipelines have mostly been developed for and validated with bacterial 16S rDNA sequences. When applying these tools on fungal ITS datasets, additional complexities (specific to ITS) have to be taken into account. First, in contrast to the near uniform length of 16S amplicons across bacterial species, the length of ITS amplicons varies substantially, between 150 and over 500 nucleotides, in different fungal species (4). Due to the current limits of Illumina-based sequencing, the longest paired-end reads measure 300 base pairs each in length, which leads to a varying overlap of the forward and reverse ITS1 amplicon reads. While the high variability in ITS sequence and length complicates the bioinformatic processing of fungal amplicon datasets, it also enables a high resolution in differentiating distinct fungal taxa (2). Second, fungal taxonomic annotation is additionally complicated by a rapidly evolving taxonomy with major reclassifications of medically relevant fungal taxa in the last few years (5). With the development of algorithms such as DADA2, which infer exact sequencing variants based on the assumed error distribution of amplicon reads, the technological requirements to discriminate sequencing variants in high resolution are available for application to ITS data sets (6). Here, we show that DADA2 can effectively discriminate ITS1 amplicons within a mock community of fungal species that are commonly identified in the human intestinal mycobiota. We further optimize the output of a DADA2-based fungal pipeline by customizing the steps from quality filtering to taxonomic annotation and applied it on patient samples.

## Results

### Denoising with DADA2 allows high-resolution discrimination of fungal ITS1 amplicon datasets

ITS1 sequences from different fungal sequences can be over 97% similar, which is beyond the resolution of operational taxonomic unit (OTU)-based classification schemes, for example UPARSE, that typically use a 3% dissimilarity threshold. To overcome this limitation, a denoising strategy like DADA2 aims to provide exact amplicon sequencing variants (ASV) that can discriminate input sequences at single nucleotide resolution (6). We compared the resolution of DADA2-computed ASVs to UPARSE-computed OTUs on a medically relevant mock fungal community that included *Aspergillus fischeri*, *Aspergillus fumigatus*, *Candida albicans*, *Candida glabrata*, *Candida metapsilosis*, two strains of *Candida parapsilosis*, *Malasezzia sympodialis*, *Meyerozyma caribbica*, *Meyerozyma guilliermondii*, and *Saccharomyces cerevisisae*.

We confirmed that DADA2 can discriminate between individual constituents that differ by a single nucleotide in ITS1 amplicons, whereas UPARSE failed to discriminate amplicon sequences with less than 3% dissimilarity (Fig. 1A and 1B). The ITS1 amplicons from *M. guilliermondii* and *M. caribbica* have 3 nucleotide differences (1% of the length), while the ITS1 amplicons of *A. fumigatus* and *A. fischeri* have a single nucleotide difference (Fig 1A). Similarly, the ITS1 amplicons of the two *C. parapsilosis* strains included in the analysis and the intragenomically heterogeneous ITS1 of *S. cerevisiae* differ by a single nucleotide (Fig 1B). Next, we wanted to confirm that DADA2 is able to differentiate these species in cases in which one species or strain is highly dominant over a species or strain with a highly similar ITS1 region. We calculated and normalized the DNA abundance of each input fungal species via quantitative PCR (“balanced”) and created extreme conditions (“extreme 1” and “extreme 2”) for the two species pairs with highly similar ITS1 amplicons *(Meyerozyma* and *Aspergillus)* by diluting one of the species 50-fold. DADA2 was able to distinguish *M. guilliermondii* and *M. caribbica* as well as *A. fumigatus* and *A. fischeri* reads even in the extreme conditions. We also demonstrated that independent UPARSE runs inconsistently assigned the sequence of the variant with most reads as the OTU reference sequence in the extreme conditions, precluding comparisons of OTU-based datasets issued from different runs (Fig. 1A).

**Figure 1.**
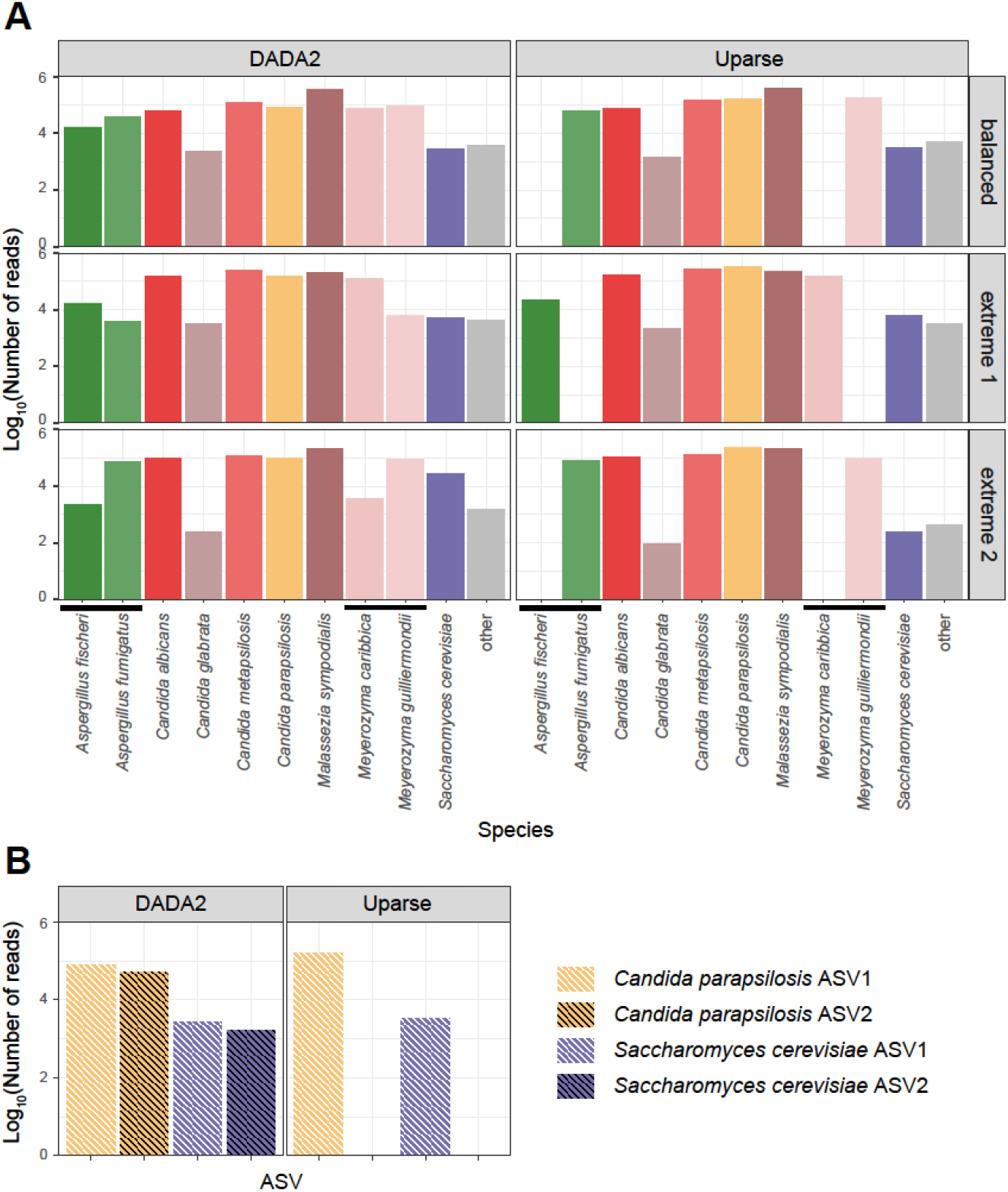
DADA2 allows a higher resolution of ITS1 sequences than UPARSE. (A) Species-level discrimination of strains in the mock community. The “balanced” community has equal 18S rDNA copy-number normalized amounts of DNA per strain. The “extreme 1” community include equal 18S rDNA copy-number normalized amounts of DNA per strain, except for *A. fumigatus* and *M. guilliermondii*, which were included at 50-fold dilution. The “extreme 2” community include equal 18S rDNA copy-number normalized amounts of DNA per strain, except for *A. fischeri* and *M. caribbica*, which were included at 50-fold dilution. (B) Sub-species level discrimination of the “balanced” community.

### Fine-tuned quality filtering variables increase the proportion of correctly denoised reads

In the DADA2 workflow, filtering is accomplished by the *filterAndTrim* function and modulated by two filtering variables: *truncQ* (truncation based on quality scores) and *maxEE* (maximum expected error). Individual reads are truncated at the first nucleotide base with a Phred quality score lower than *truncQ*. After truncation, reads with a number of expected errors equal to or higher than *maxEE* are removed. The default value for both *truncQ* and *maxEE* is 2 in the DADA2 package. In the initial manuscript introducing the concept of *maxEE* the authors suggested a value of 1, corresponding to no expected error (7). Since different filtering strategies have not been evaluated for fungal datasets, we tested different combinations of *truncQ* and *maxEE*, using values between 1 and 10 for both filtering variables. We compared the dereplicated and denoised output of these 100 runs to the expected sequences within our balanced mock community. The overall number of reads and the number of expected reads returned by the pipeline increased when both *maxEE* and *truncQ* values were increased (Fig. 2A). The proportion of noise (ratio of non-expected to expected) reads decreased primarily with an increase in the *truncQ* value. A value of 8 for both *truncQ* and *maxEE* maximized the number and proportion of correctly identified sequences in the dataset.

**Figure 2.**
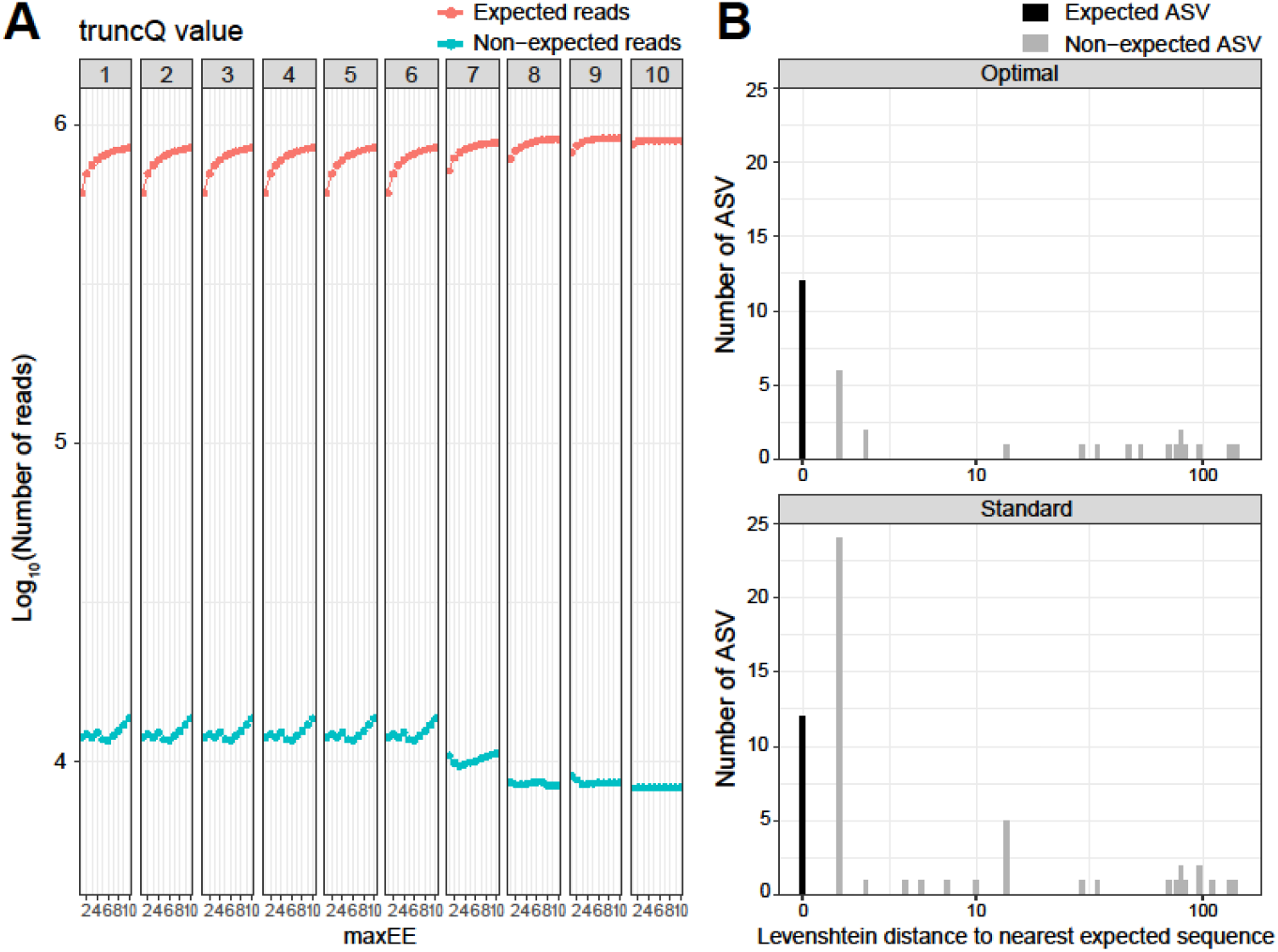
Impact of varying the filtering variables truncQ and maxEE on the output of DADA2. (A) Ratio of expected to non-expected sequences (based on the strains within the mock community) for different combinations of truncQ and maxEE. (B) Comparison of the Levenshtein distance for all ASV reference sequences to the nearest expected read between standard values (maxEE=2, truncQ=2) and optimized values (maxEE=8, truncQ=8) of the filtering variables.

With this optimal filtering combination, the number of expected reads increased from 700,747 (default filter values) to 897,992. The number of non-expected reads decreased from 12,207 to 8,428. With both the optimal and default filtering combinations, we retrieved all 12 expected ASVs. The number of non-expected ASVs (noise) decreased from 46 to 22 when the optimal filtering values were used. The decrease in non-expected ASVs is mainly due to ASVs that differed from the expected sequence by 14 nucleotides or less (Fig. 2B).

### Species-specific bias in ITS-based amplicon pipelines

We detected species-specific filtering biases that are independent of amplicon length: with standard filtering values used in the *filterandTrim* function, a majority of reads pertaining to *Aspergillus* species would have been discarded, and to a lesser extent reads pertaining to *S. cerevisiae* and *C. glabrata* (Fig 3A). Increasing *maxEE* and *truncQ* values maintained a higher number of reads pertaining to these species. We confirmed species-specific differences in the quality of Illumina reads with the highest median expected error for *Aspergillus* reads compared to reads from other species (Table 1, Figs 3B, S1, and S2).

**Figure 3.**
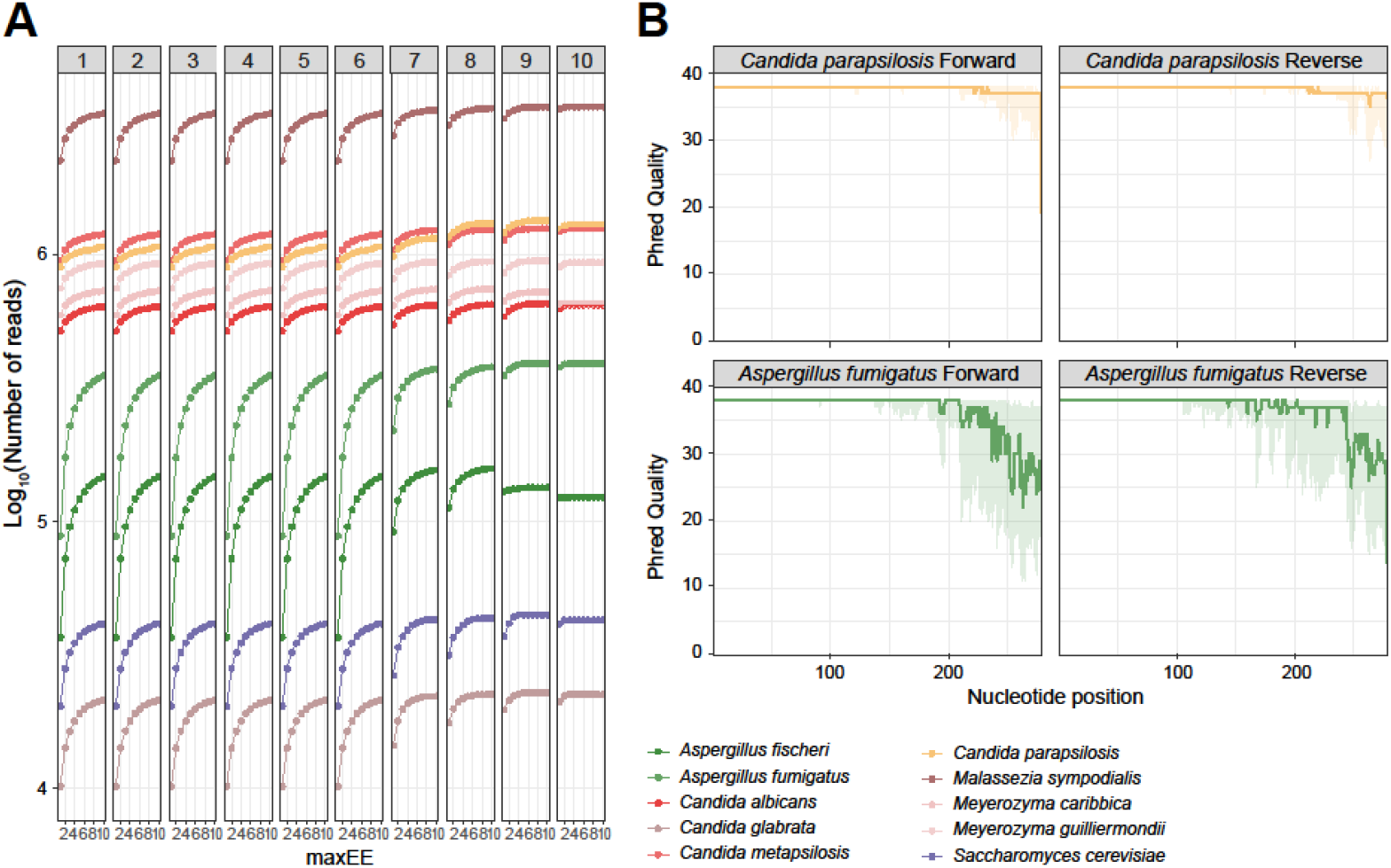
Species-specific biases in fungal ITS1 amplicon sequencing and analysis. (A) Impact of varying truncQ and maxEE on the number of species-specific reads. (B) Representative quality profile of raw reads that were denoised into exact sequence matches to *A. fumigatus* and *C. albicans*. The line represents the median Phred Score at that position, while the shaded area represents the 25^th^ to 75^th^ percentiles.

**Table 1:** Expected errors (EE) of species-specific forward and reverse reads

| Forward reads |  | Reverse reads |  |
| --- | --- | --- | --- |
| Taxon | Median<br>EE | Taxon | Median EE |
| A. fumigatus | 0.89 | A. fumigatus | 0.82 |
| A. fischeri | 0.86 | A. fischeri | 0.87 |
| C. albicans | 0.06 | C. albicans | 0.07 |
| C. glabrata | 0.37 | C. glabrata | 0.42 |
| C. metapsilosis | 0.08 | C. metapsilosis | 0.12 |
| C. parapsilosis | 0.06 | C. parapsilosis | 0.11 |
| M. sympodialis | 0.23 | M. sympodialis | 0.14 |
| M. caribbica | 0.07 | M. caribbica | 0.08 |
| M.<br>guilliermondii | 0.07 | M. guilliermondii | 0.08 |
| S. cerevisiae | 0.15 | S. cerevisiae | 0.61 |

Next, we assessed the effect of customizing the filtering strategy on patient fecal samples. (Fig 4A). We could confirm that the customization enables us to significantly increase the relative abundance of reads pertaining to *Aspergillus* species (Samples A-C). By using the standard filtering variables, no *Aspergillus* would have been detected in Sample C. To a lesser extent, the relative abundance of reads pertaining to *S. cerevisiae* could also been increased (Samples D-F). The increase in relative abundance of these species correlated with an increase in the total number of reads that were retained by using the optimized filtering strategy (Fig. 4B).

**Figure 4:**
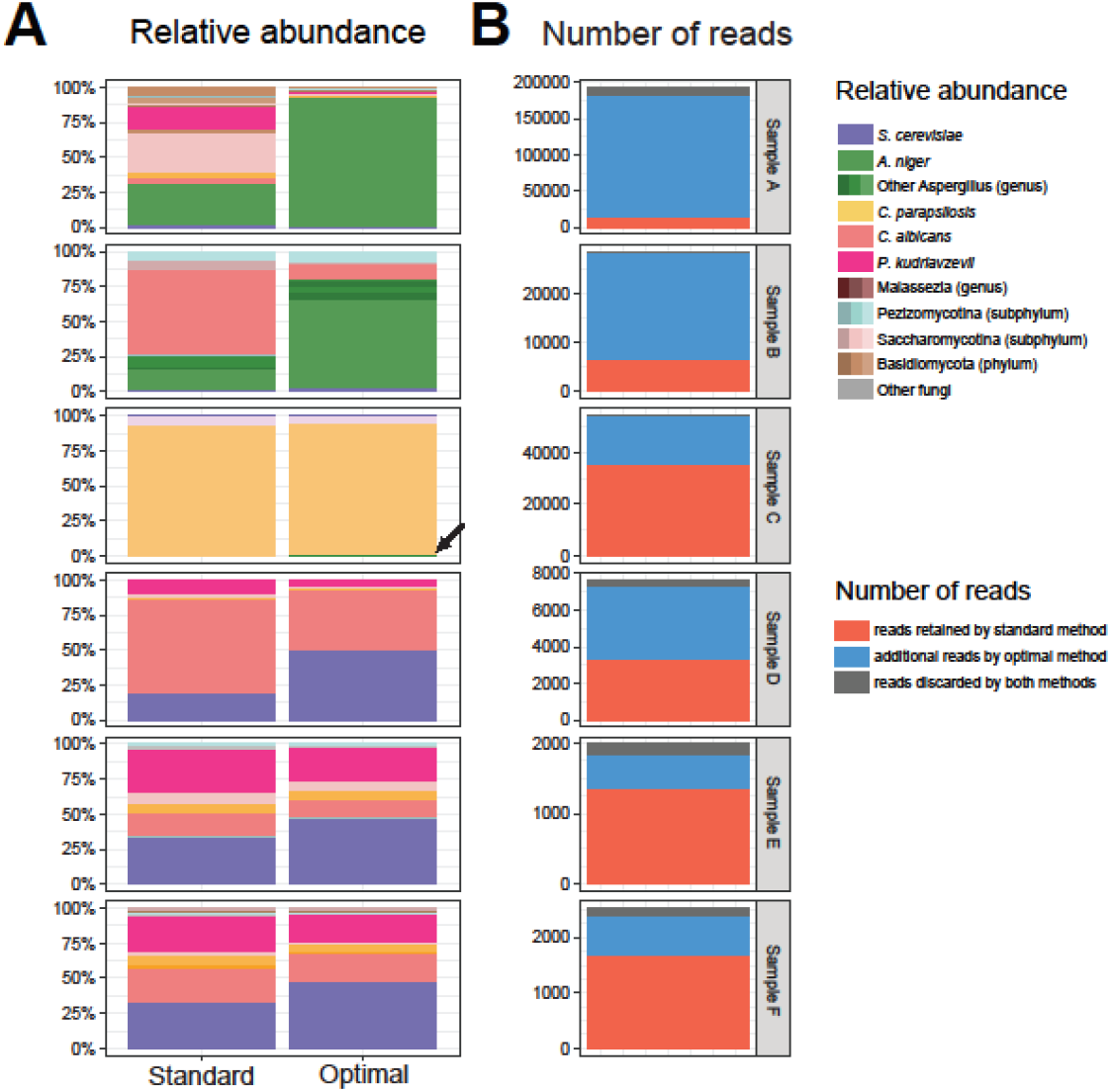
Effect of customizing filtering values on fecal samples. (A) Taxonomic composition of fecal samples according to the filtering strategy used. The arrow shows the retention of reads from Aspergillus species which were completely discarded by the standard filtering strategy. (B) Number of reads retained by DADA2 according to filtering strategy used.

Although we normalized the amount of input rDNA in the “balanced” sample and adapted the filtering parameters, the number of reads attributed to *C. glabrata* and *S. cerevisiae* were about a log lower than reads of other species (Fig. 1A) (8). This is explained in part by sequencing bias from the Illumina platform against the longer ITS1 amplicon of these two species (Fig. 5A). Additionally, we hypothesized that a single nucleotide difference in the primer region of the forward primer (ITS1-F) between the reference genome of *S.cerevisiae* and of some strains of *C. glabrata*, and other fungal taxa could be responsible for a portion of the observed species-specific bias. (Fig 5B). By using an alternative primer with a wobble nucleotide at the diverging position near the 3, end, the yield of *C. glabrata* and *S. cerevisiae* reads could be increased 7.2-fold in the balanced mock community and 2.0-fold in a community maximally enriched in both species (90% of input DNA, Fig. 5C).

**Figure 5.**
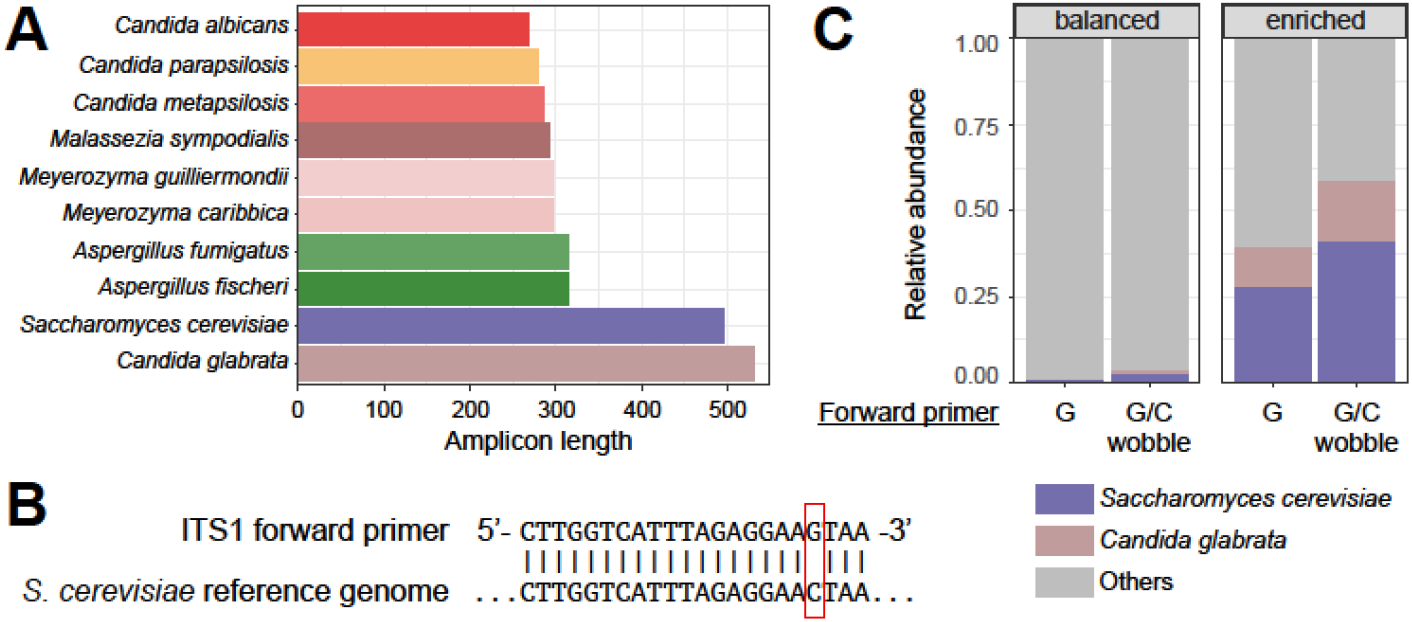
Length-specific biases in fungal ITS1 amplicon sequencing and adaptation of forward primers for better recall of *S. cerevisiae* and *C. glabrata*. (A) Length variation in the amplified ITS1 region of strains within the mock community. (B) Single nucleotide difference between the ITS1-F primer and the *S. cerevisiae* reference genome. (C) Impact on the relative abundance of *S. cerevisiae* and *C. glabrata* when using a wobble forward primer allowing for the single nucleotide difference between the ITS1-F primer and the *S. cerevisiae* reference genome.

### Taxonomic annotation for fungal ITS1 amplicon datasets

Multiple combinations of annotation algorithms and reference databases have been developed for taxonomic annotation. To find an optimal combination, we compared three regularly updated reference databases (UNITE with three different versions, NCBI NT, and NCBI ITS RefSeq Fungi) and two annotation algorithms (RDP and BLAST). DADA2 implements a naïve Bayes classifier (RDP classifier) in its *assignTaxonomy* function. The ITS-specific workflow recommends using a UNITE database without specifying which database version to use (https://benjjneb.github.io/dada2/ITS_workflow.html). We tested three different versions of the UNITE database: 1) including singletons as reference sequences, 2) including global and 97% singletons, 3) the full UNITE+INSD database. All three databases resulted in the correct taxonomic annotation at the genus level. With the default bootstrap threshold of 50, species-level annotation was not achieved in four or five of 12 ASVs, depending on the database used. (Table 2). With the bootstrap threshold set at 80 (suggested for reads longer than 250nts (9)), the proportion of species-level annotation decreased further. Species-level identification was inconsistent between the different versions of the UNITE database (Table 2). Due to the random nature of the naïve Bayes classifier algorithm, identification at the species was not reproducible between different runs (Table S2).

**Table 2:** Comparison of taxonomic annotation at the Genus and Species level by three versions of the UNITE database and the *assignTaxonomy* function of dada2 by using a seed of 100. UNITE_s: UNITE database including global and 97% singletons; UNITE: UNITE database including singletons set as reference sequences; UNITE+INSD: full UNITE and INSD database. * Top hit is a sequence without Species-level annotation in UNITE. ** Top hit is a sequence with an undefined Species-level annotation in UNITE.

| Expected | UNITE_s |  |  |  | UNITE |  |  |  | UNITE+INSD |  |  |  |
| --- | --- | --- | --- | --- | --- | --- | --- | --- | --- | --- | --- | --- |
|  | Genus | Bootstrap | Species | Bootstrap | Genus | Bootstrap | Species | Bootstrap | Genus | Bootstrap | Species | Bootstrap |
| A. fischeri | Aspergillus | 99 | NA | 18 | Aspergillus | 100 | A. fischeri | 50 | Aspergillus | 100 | NA | 48 |
| A. fumigatus | Aspergillus | 100 | NA | 40 | Aspergillus | 100 | NA | 23 | Aspergillus | 100 | A. fumigatus | 76 |
| C. albicans | Candida | 100 | C. albicans | 93 | Candida | 100 | C. albicans | 99 | Candida | 100 | C. albicans | 74 |
| C. glabrata | Nakaseomyces | 100 | NA* | 100 | Nakaseomyces | 100 | NA* | 100 | Nakaseomyces | 95 | C. glabrata | 95 |
| C. metapsilosis | Candida | 99 | C. metapsilosis | 86 | Candida | 100 | C. metapsilosis | 99 | Candida | 100 | C. metapsilosis | 100 |
| C. parapsilosis | Candida | 100 | C. parapsilosis | 99 | Candida | 100 | NA | 46 | Candida | 100 | C. parapsilosis | 100 |
| C. parapsilosis | Candida | 100 | C. parapsilosis | 99 | Candida | 100 | C. parapsilosis | 55 | Candida | 100 | C. parapsilosis | 100 |
| M. carribica | Meyerozyma | 100 | M. guilliermondii | 97 | Meyerozyma | 100 | M. guilliermondii | 92 | Meyerozyma | 100 | M. carpophila | 96 |
| M. guilliermondii | Meyerozyma | 100 | M. guilliermondii | 99 | Meyerozyma | 100 | M. guilliermondii | 100 | Meyerozyma | 100 | NA | 46 |
| M. sympodialis | Malassezia | 100 | M. sympodialis | 100 | Malassezia | 100 | M. sympodialis | 100 | Malassezia | 100 | M. sympodialis | 100 |
| S. cerevisiae | Saccharomyces | 100 | S. cerevisiae | 87 | Saccharomyces | 100 | S. cerevisiae | 89 | Saccharomyces | 100 | Saccharomyces species** | 100 |
| S. cerevisiae | Saccharomyces | 100 | S. cerevisiae | 95 | Saccharomyces | 100 | S. cerevisiae | 92 | Saccharomyces | 100 | Saccharomyces species** | 100 |

In contrast to the RDP classifier, BLAST-based algorithms do not return a single, most likely hit, but rather a list of the top potential hits based on E-value or other scores (full output in Supplementary material), which is representative of the inherent uncertainties in taxonomic annotation. We arranged these ties based on whether a species-level designation was available and based on the number of times a specific species-level designation was returned. With this strategy, both the full UNITE+INSD and the NCBI NT databases allowed a correct species-level annotation for all query sequences (Tables 3 and 4). In contrast, the BLAST-based algorithm was ineffective in returning a correct species-level annotation with the alternative novel fungal database NCBI ITS_RefSeq_Fungi (Table S3).

**Table 3:**
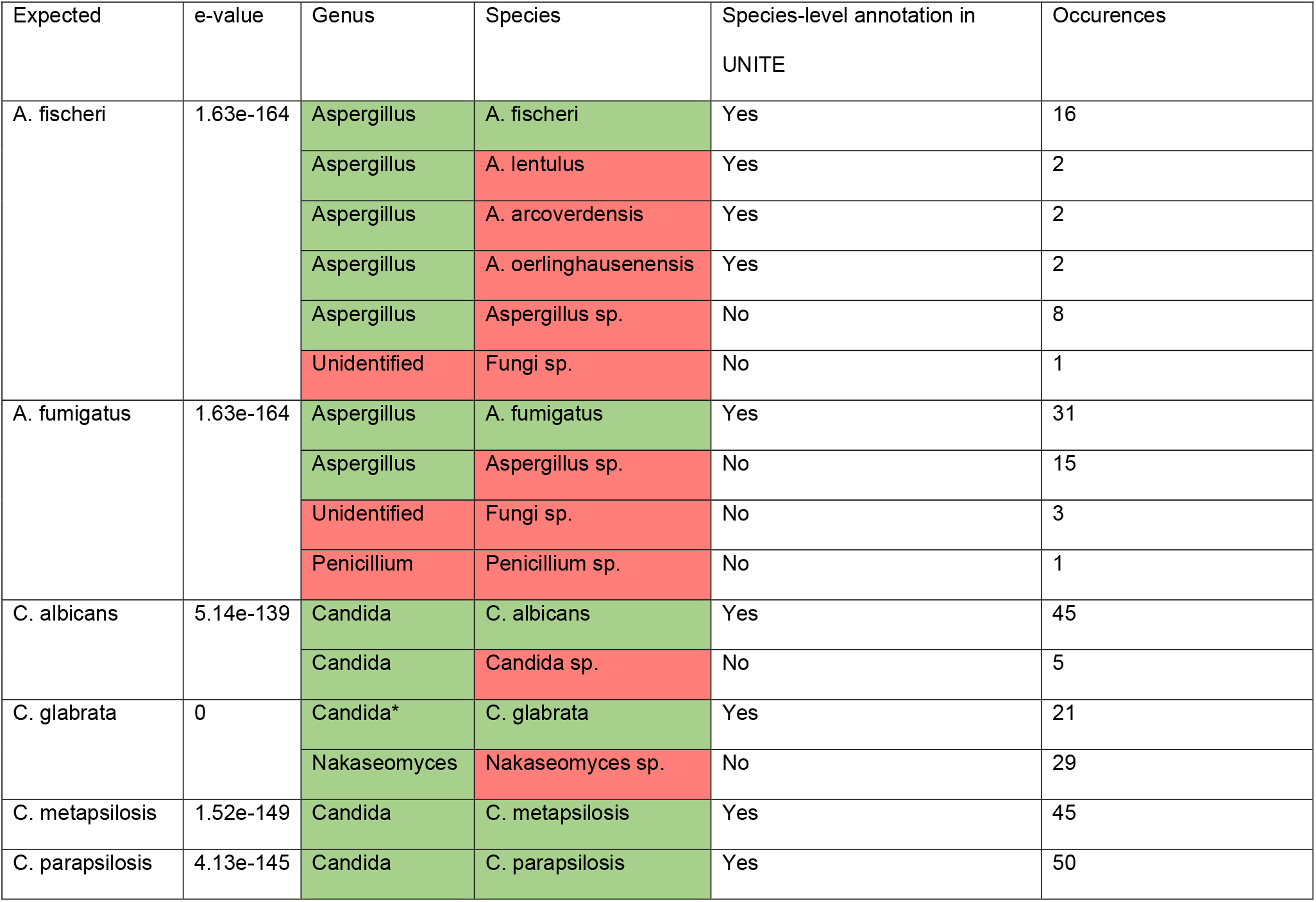
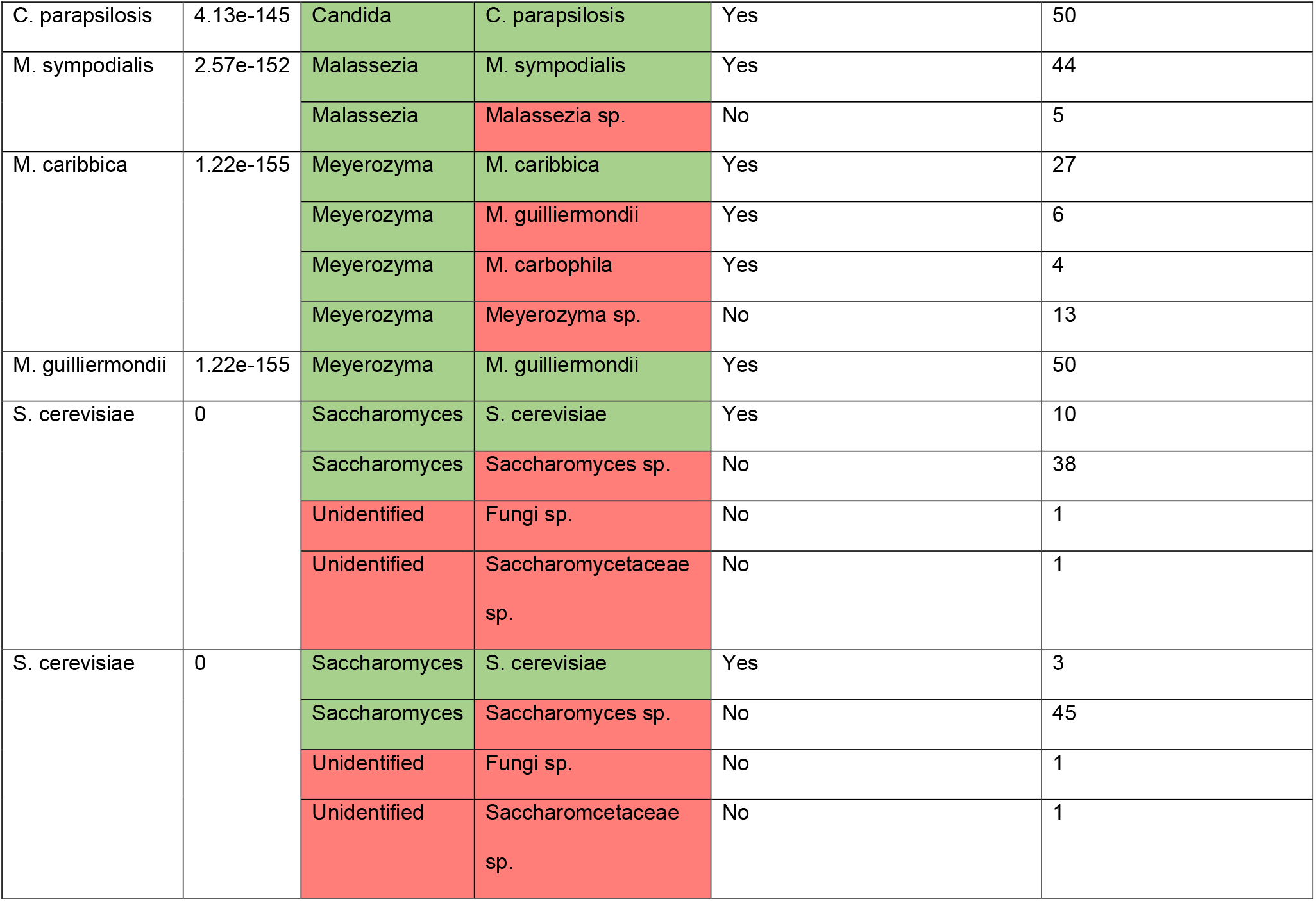
Tied top hits with lowest e-value returned from a BLAST-based algorithm with the UNITE+INSD database as training set. Occurrences: number of times the same taxonomic annotation was returned. * Taxonomy in the UNITE database has not yet been updated to reflect new nomenclature (Nakaseomyces).

**Table 4:** Tied top hits with lowest e-value returned from a BLAST-based algorithm with the NT database as training set. Only returned hits with a t least a sequence-based annotation are shown. Occurrences: number of times the same taxonomic annotation was returned.

| Expected | e-value | Genus | Species | Occurences |
| --- | --- | --- | --- | --- |
| A. fischeri | 1.24e-161 | Aspergillus | A. fischeri | 11 |
|  |  | Aspergillus | A. lentulus | 10 |
|  |  | Aspergillus | A. fumigatiaffinis | 5 |
|  |  | Aspergillus | A. arcoverdensis | 3 |
|  |  | Aspergillus | A. oerlinghausensis | 3 |
|  |  | Aspergillus | Aspergillus sp. | 2 |
|  |  | Aspergillus | A. novofumigatus | 1 |
| A. fumigatus | 1.24e-161 | Aspergillus | A. fumigatus | 49 |
|  |  | Aspergillus | Aspergillus sp. | 1 |
| C. albicans | 3.87e-136 | Candida | C. albicans | 45 |
|  |  | Candida | Candida sp. | 5 |
| C. metapsilosis | 1.15e-146 | Candida | C. metapsilosis | 8 |
| C. parapsilosis | 3.11e-142 | Candida | C. parapsilosis | 30 |
| C. parapsilosis | 3.11e-142 | Candida | C. parapsilosis | 49 |
| M. sympodialis | 1.94e-149 | Malassezia | M. sympodialis | 19 |
|  |  | Malassezia | M. globosa. | 5 |
| M. caribbica | 9.19e-153 | Meyerozyma | M. caribbica | 24 |
|  |  | Meyerozyma | M. guilliermondii | 3 |
|  |  | Meyerozyma | M. carpophila | 2 |
|  |  | Meyerozyma | Meyerozyma sp. | 1 |
|  |  | Candida | Candida sp. | 1 |
| M. guilliermondii | 9.19e-153 | Meyerozyma | M. guilliermondii | 48 |
|  |  | Meyerozyma | M. carpophila | 1 |
|  |  | Saccharomyces | Saccharomyces sp. | 1 |
| S. cerevisiae | 0 | Saccharomyces | S. cerevisiae | 1024 |
|  |  | Saccharomyces | S. cf<br>cerevisiae/paradoxus | 1 |
| S. cerevisiae | 0 | Saccharomyces | S. cerevisiae | 212 |

## Discussion

This study demonstrated that a DADA2-based denoising algorithm distinguished fungal ITS1 amplicon reads that differ by a single nucleotide, as previously demonstrated for bacterial 16S amplicon datasets (6). This discriminatory power allowed for species-level distinctions for the members in our mock community of medically relevant fungi. This discriminatory power was not achievable by an OTU-based approaches due to grouping clusters of sequences with 97% similarity (10). Additionally, independent DADA2 runs yield the same ASV classification, allowing comparisons between different studies.

The high resolution provided the possibility to identify intraspecies variability, provided that there is a difference in the ITS1 amplicon, as shown in the discrimination of two *C. parapsilosis* strains with a single nucleotide polymorphism. This fine discriminatory power was harnessed to track individual *C. parapsilosis* strains across different body sites and time to determine the relationship of intestinal and bloodstream isolates in a pathogenesis study (11). In *S. cerevisiae* and certain other species, fungal rDNA is present in multiple copies and can contain intragenomic polymorphisms, resulting in the presence of more than one ASV for a given clone (12, 13). The optimized DADA2 pipeline can discriminate these polymorphisms and return distinct ASVs for a single clonal origin. On the other end of the spectrum, it is possible that two fungal species have an identical ITS1 (14). Thus, it is important to note that fungal ITS1-based ASVs are not a substitute to define a specific fungal species and diversity measures may be overestimated at the fungal ASV level (15). We feel these limitations are clearly outweighed by the benefit of higher taxonomic resolution associated with an ASV-based approach compared to an OTU-based approach. If needed, clustering at a higher (taxonomy-based) level is still possible for analytic purposes.

Due to the decrease in sequencing quality towards the end of Illumina reads, it is generally recommended to trim reads at the 3’ end in processing bacterial 16S data (6). With this trimming step, the overall quality increases and more reads can pass the quality filter implemented in the pipeline (16). With the variation in ITS amplicon length, this approach is not generally recommended for fungal datasets. Besides the possibility of trimming reads at fixed length, the *filterandTrim* function of the DADA2 package incorporates the possibility to trim reads at the first position with a Phred score lower than a prespecified threshold. By default, this threshold is set to 2. In this study, we showed that increasing this threshold to 8 increased the number of reads that passed the second quality filter step and were denoised correctly.

Quality filtering with the *filterandTrim* function was performed by removing reads with a higher expected error than a specified threshold. Using the number of expected errors within a read has been shown to be a superior filtering strategy to using the overall or average quality of a read (7). In our dataset, we showed that increasing the threshold leads to a better recovery of correct reads. Since DADA2 relies on the distribution of sequencing errors, we speculate that including a higher number of erroneous reads may increase the reliability of the error model. Intriguingly, changing the filtering and trimming parameters affected specific fungal taxa differentially, a phenomenon that has not been described widely in the literature. The ITS regions of different fungal taxa vary considerably in sequence, length, and GC content (17). The impact on differential read quality has to be considered in the analysis of ITS datasets to minimize this taxon-specific filtering bias.

An additional species-specific bias can be introduced by the commonly used ITS1 forward primer, as it differs by a single nucleotide from the complementary regions for taxa such as *S. cerevisiae* and *C. glabrata*. The impact of this primer modification is measurable, yet moderate in comparison to other biases, such as variations in amplicon length and in rDNA copy numbers between taxa (18). While ITS-based mycobiota analysis will detect different members of fungal communities in high resolution, it can only approximate their relative abundance. However, it remains an extremely valuable tool to classify community members, to assess temporal and spatial variations of the mycobiota, and to monitor exponential expansion of pathogenic fungal taxa seen in specific disease states (3).

Shotgun metagenomics may provide a less biased analysis of microbial communities. However, in most communities, such as the human intestine, the overall abundance of fungi is low.

Without the enrichment step inherent to amplicon sequencing, these fungal communities cannot be readily detected in shotgun metagenomic datasets at current sequencing depths. In addition, reference databases for shotgun metagenomic analyses are either absent or incomplete (19). At the present time, ITS-based amplicon approaches remain a cost-efficient standard to profile fungal communities.

Correct genus-level assignment was achievable for ITS1 amplicon datasets either via the RDP naïve Bayesian classifier implemented in DADA2 or via a BLAST-based approach, irrespective of the reference database. However, both approaches had limitations. The RDP classifier returned a single hit or no hit if the bootstrap value is below a prespecified threshold, i.e. the assignment is ambiguous (20). Additionally, the randomness inherent to the RDP classifier led to varying results between different runs. Slight differences in the version of the database gave rise to inconsistent species-level results. Overall, in our mock community, the RDP algorithm was suboptimal for species-level annotation. A BLAST-based approach is limited by the fact that the single best hit is obligately returned, since the algorithms stops after a certain number of hits (that can be customized) have passed the e-value threshold. The BLAST-based approach has the advantage of listing several potential hits (in order of likelihood), illustrating the uncertainty of taxon calling, and of narrowing down the possible correct assignment. To confirm biologically meaningful associations based on mycobiota data, it is desirable to confirm taxonomic annotations by culture-based methods.

In this study, we achieved similar levels of species-level annotation in our BLAST-based algorithm with both databases NCBI NT, and the full UNITE+INSD. Using the NCBI ITS RefSeq database did not result in correct taxonomic annotation at the species level. Of note, the downloadable UNITE databases are updated once yearly while NCBI databases and the linked NCBI taxonomy are updated continuously (21, 22). In the 2020 iteration of UNITE used for this study, the taxonomy for *Candida* strains was not yet updated to reflect the family/genus denominations (*Debaryomycetaceae* as the family for *C. albicans*, *C. parapsilosis*, and *C. metapsilosis*, and *Saccharomycetaceae* as family and *Nakaseomyces* as genus for *C. glabrata*). It is important to use the newest version of either database to reflect the rapidly changing fungal taxonomy (5, 21).

In summary, our mock community was representative of commonly seen medically relevant fungi and we established that a DADA2 based pipeline can discriminate ITS1 amplicons with single nucleotide resolution as a proof-of-concept. While ITS-inherent species-specific biases cannot be overcome fully, the DADA2-based analytic pipeline can be optimized by customizing quality trimming and filtering, and taxonomic annotation.

## Methods

### Fungal strains and DNA preparation

We selected 11 different fungal strains from 10 distinct species for analysis (Table S4). These strains were chosen to reflect a range of distinct medically relevant fungi and included strains with more than 97% identity in the ITS1 amplicon (*A. fumigatus* and *A. fischeri*, *M. caribbica* and *M. guilliermondii*, and two *C. parapsilosis* strains). Fungal strains were revived from glycerol stock and streaked on YPD agar, cultured at 37°C for overnight. Then the strains were inoculated in YPD liquid medium and cultured at 37°C, 240rpm for overnight. Fungal cells were harvested and washed twice with sterile water. Fungal DNA was extracted with the QIAamp DNA mini kit (Qiagen 51306).

### Composition of DNA pools

The 18S copy number per μl of DNA for each strain was measured by quantitative PCR (23). DNA of all the strains was pooled at equal amount of 18S copy numbers for the “balanced” community. For the Extreme 1 community, equal amounts of DNA were pooled for all strains except for *A. fumigatus* and *M. guilliermondii*, which were both diluted 50-fold. For the Extreme 2 community equal amounts of DNA were used for all strains except for *A. fischeri* and *M. caribbica*, which were both diluted 50-fold. Finally, the Enriched community was composed of 10% of the balanced community, 45% of *S. cerevisiae*, and 45% *C. glabrata* DNA.

### Fecal samples

Fecal samples were drawn from a fecal biorepository of patients undergoing allogeneic hematopoietic cell transplantation at Memorial Sloan Kettering Cancer Center (11, 24). We selected fecal samples from different sequencing runs and that contained reads attributed to *Aspergillus*. Samples were processed as described previously(11).

### Amplicon production and sequencing

We amplified the ITS1 region with the primer set ITS-1-F (5’-CTTGGTCATTTAGAGGAAGTAA-3’) and 5.8S-1R (5’-GTTCAAAGAYTCGATGATTCAC-3’). We also tested an alternative forward primer including a replacement wobble nucleotide (5’-CTTGGTCATTTAGAGGAA<u>**S**</u>TAA-3’). The DNA was amplified for 35 cycles (98° C, 53° C, and 72° C, for 30 s each) using phusion polymerase (F530L), as reported previously (11). The ensuing amplicons were sequenced on an Illumina Miseq platform with paired-end 300 setting. The amplicon and sequencing strategy results in both forward and reverse reads being present in the R1 and R2 reads. The raw reads were preprocessed by separating forward and reverse reads based on primer presence into two different files. Subsequently primers and (partial) read-ins into the opposite primer were removed by using *cutadapt* (25).

### Denoising and OTU clustering

Denoising was performed using the DADA2 package in R (6). No fixed length trimming was used. To test different filtering strategies 100 iterations of the *filterandTrim* function with maxEE and truncQ values varying between 1 and 10 each were performed on the preprocessed reads of the “balanced” community with the ITS-1-F/5.8S-1R primer set. For all other analyses, the ASV object obtained by using *maxEE* and *truncQ* of 8 each was used. OTU clustering was performed via UPARSE by using a customized pipeline based on USEARCH and VSEARCH using the suggested value of *maxEE* of 1(26–28).

### Taxonomic annotation

To test the RDP Naïve Bayes classifier implemented in the *assignTaxonomy* function of DADA2, we downloaded three variants of the UNITE database, version 8.2 (February 2020) (29). DADA2 can utilize two variants of the general FASTA release, one that includes singletons as reference sequences (DOI: 10.15156/BIO/786368) and another that includes global and 97% singletons (DOI: 10.15156/BIO/786368). The third variant consists of the full UNITE and INSD dataset (DOI: 10.15156/BIO/786372). The header of this dataset was reformatted to comply with DADA2 requirements. We used a default DADA2 bootstrap threshold of 50.

To test a BLAST-based approach to taxonomic assignment, the UNITE and INSD dataset was converted to a BLAST database, NCBI NT and NCBI RefSeq ITS libraries were downloaded in December 2020 from the NCBI FTP site (21, 28, 30). We performed a local BLAST search for the expected sequences with a maximum of 50 target sequences. We calculated the number of times a specific species-level taxonomy was returned per sequence for the NT and the UNITE databases. This was not possible due to the nature of the NCBI ITS database which includes a unique sequence per species. Additionally, for the UNITE database we sorted the results on whether a species-level annotation was available or not. The full BLAST output is available in the Supplementary material.

### Analysis

All analyses were performed using R version 3.6.3 (The R Foundation for Statistical Computing, Vienna, Austria).

### Data availability

The preprocessed reads and the code will be available on Github after acceptance of the manuscript.

### Study approval

Patients provided written informed consent for biospecimen collection. The fecal biospecimen repository was approved by the MSKCC institutional review board. Is

## Author contributions

TR, BZ, TMH, and YT conceived the study. TR and BZ handled sample processing, DNA extraction, and amplicon preparation. TR analyzed the data with assistance by JF and YT. TR and TMH wrote the first draft of the manuscript with subsequent contributions by all coauthors. All authors approved the submitted version of the manuscript.

## Acknowledgments

This work was supported by Deutsche Forschungsgemeinschaft (DFG, German Research Foundation) grant RO-5328/2 (T.R.), National Institutes of Health (NIH) grants R01 AI093808 (T.M.H.), R21 AI105617 (T.M.H.), R21 AI156157 (T.M.H.), R01 AI137269 (Y.T.) and NIH P30 CA008748 (Cancer Center Core Grant).

## References

1. Lynch SV, and Pedersen O. The Human Intestinal Microbiome in Health and Disease. N Engl J Med. 2016;375(24):2369–79.

2. Schoch CL, Seifert KA, Huhndorf S, Robert V, Spouge JL, Levesque CA, et al. Nuclear ribosomal internal transcribed spacer (ITS) region as a universal DNA barcode marker for Fungi. Proc Natl Acad Sci U S A. 2012;109(16):6241–6.

3. Zhai B, Ola M, Rolling T, Tosini NL, Joshowitz S, Littmann ER, et al. High-resolution mycobiota analysis reveals dynamic intestinal translocation preceding invasive candidiasis. Nat Med. 2020.

4. Motooka D, Fujimoto K, Tanaka R, Yaguchi T, Gotoh K, Maeda Y, et al. Fungal ITS1 Deep-Sequencing Strategies to Reconstruct the Composition of a 26-Species Community and Evaluation of the Gut Mycobiota of Healthy Japanese Individuals. Front Microbiol. 2017;8:238.

5. Borman AM, and Johnson EM. Name changes for fungi of medical importance, 2018-2019. J Clin Microbiol. 2020.

6. Callahan BJ, McMurdie PJ, Rosen MJ, Han AW, Johnson AJ, and Holmes SP. DADA2: High-resolution sample inference from Illumina amplicon data. Nat Methods. 2016;13(7):581–3.

7. Edgar RC, and Flyvbjerg H. Error filtering, pair assembly and error correction for next-generation sequencing reads. Bioinformatics. 2015;31(21):3476–82.

8. Gohl DM, Magli A, Garbe J, Becker A, Johnson DM, Anderson S, et al. Measuring sequencer size bias using REcount: a novel method for highly accurate Illumina sequencing-based quantification. Genome Biol. 2019;20(1):85.

9. Edgar RC. Accuracy of taxonomy prediction for 16S rRNA and fungal ITS sequences. PeerJ. 2018;6:e4652.

10. Callahan BJ, McMurdie PJ, and Holmes SP. Exact sequence variants should replace operational taxonomic units in marker-gene data analysis. ISME J. 2017;11(12):2639–43.

11. Zhai B, Ola M, Rolling T, Tosini NL, Joshowitz S, Littmann ER, et al. High-resolution mycobiota analysis reveals dynamic intestinal translocation preceding invasive candidiasis. Nat Med. 2020;26(1):59–64.

12. Zhao Y, Tsang CC, Xiao M, Cheng J, Xu Y, Lau SK, et al. Intra-Genomic Internal Transcribed Spacer Region Sequence Heterogeneity and Molecular Diagnosis in Clinical Microbiology. Int J Mol Sci. 2015;16(10):25067–79.

13. Simon UK, and Weiss M. Intragenomic variation of fungal ribosomal genes is higher than previously thought. Mol Biol Evol. 2008;25(11):2251–4.

14. Seifert KA, Samson RA, Dewaard JR, Houbraken J, Levesque CA, Moncalvo JM, et al. Prospects for fungus identification using CO1 DNA barcodes, with Penicillium as a test case. Proc Natl Acad Sci U S A. 2007;104(10):3901–6.

15. Lena FEE, Maurice S, Morgado L, Martin-Sanchez PM, Skrede I, and Kauserud H. The influence of intraspecific sequence variation during DNA metabarcoding: A case study of eleven fungal species. Mol Ecol Resour. 2021.

16. Mohsen A, Park J, Chen YA, Kawashima H, and Mizuguchi K. Impact of quality trimming on the efficiency of reads joining and diversity analysis of Illumina paired-end reads in the context of QIIME1 and QIIME2 microbiome analysis frameworks. BMC Bioinformatics. 2019;20(1):581.

17. Yang RH, Su JH, Shang JJ, Wu YY, Li Y, Bao DP, et al. Evaluation of the ribosomal DNA internal transcribed spacer (ITS), specifically ITS1 and ITS2, for the analysis of fungal diversity by deep sequencing. PLoS One. 2018;13(10):e0206428.

18. Lofgren LA, Uehling JK, Branco S, Bruns TD, Martin F, and Kennedy PG. Genome-based estimates of fungal rDNA copy number variation across phylogenetic scales and ecological lifestyles. Mol Ecol. 2019;28(4):721–30.

19. Rolling T, Hohl TM, and Zhai B. Minority report: the intestinal mycobiota in systemic infections. Curr Opin Microbiol. 2020;56:1–6.

20. Wang Q, Garrity GM, Tiedje JM, and Cole JR. Naive Bayesian classifier for rapid assignment of rRNA sequences into the new bacterial taxonomy. Appl Environ Microbiol. 2007;73(16):5261–7.

21. Robbertse B, Strope PK, Chaverri P, Gazis R, Ciufo S, Domrachev M, et al. Improving taxonomic accuracy for fungi in public sequence databases: applying ‘one name one species’ in well-defined genera with Trichoderma/Hypocrea as a test case. Database (Oxford). 2017;2017.

22. Schoch CL, Ciufo S, Domrachev M, Hotton CL, Kannan S, Khovanskaya R, et al. NCBI Taxonomy: a comprehensive update on curation, resources and tools. Database (Oxford). 2020;2020.

23. Liu CM, Kachur S, Dwan MG, Abraham AG, Aziz M, Hsueh PR, et al. FungiQuant: a broad-coverage fungal quantitative real-time PCR assay. BMC Microbiol. 2012;12:255.

24. Peled JU, Gomes ALC, Devlin SM, Littmann ER, Taur Y, Sung AD, et al. Microbiota as Predictor of Mortality in Allogeneic Hematopoietic-Cell Transplantation. N Engl J Med. 2020;382(9):822–34.

25. Martin M. Cutadapt removes adapter sequences from high-throughput sequencing reads. EMBnet journal. 2011;17(1):10–2.

26. Edgar RC. UPARSE: highly accurate OTU sequences from microbial amplicon reads. Nat Methods. 2013;10(10):996–8.

27. Edgar RC. Search and clustering orders of magnitude faster than BLAST. Bioinformatics. 2010;26(19):2460–1.

28. Rognes T, Flouri T, Nichols B, Quince C, and Mahe F. VSEARCH: a versatile open source tool for metagenomics. PeerJ. 2016;4:e2584.

29. Nilsson RH, Larsson KH, Taylor AFS, Bengtsson-Palme J, Jeppesen TS, Schigel D, et al. The UNITE database for molecular identification of fungi: handling dark taxa and parallel taxonomic classifications. Nucleic Acids Res. 2019;47(D1):D259–D64.

30. Clark K, Karsch-Mizrachi I, Lipman DJ, Ostell J, and Sayers EW. GenBank. Nucleic Acids Res. 2016;44(D1):D67–72.

