## Supplemental File 1 for "Customization of a *dada2*-based pipeline for fungal Internal Transcribed Spacer 1 (ITS 1) amplicon datasets"

| **Species / Strain** | **Reference** |
|---|---|
| Candida albicans SC5314 | (31) |
| Candida parapsilosis ASV1 | Clinical strain from Memorial Sloan Kettering Cancer Center (MSKCC) (11) |
| Candida parapsilosis ASV2 | Clinical strain from MSKCC (11) |
| Meyerozyma caribbica | ATCC 28873™ |
| Meyerozyma guillermondii | ATCC 6260™ |
| Aspergillus fumigatus AF293 | (32) |
| Apergillus fischeri | ATCC 1020™ |
| Saccharomyces cerevisiae | Clinical strain from MSKCC |
| Candida metapsilosis | Clinical strain from MSKCC (11) |
| Malassezia sympodialis | (33) |
| Candida glabrata | Clinical strain from MSKCC |

Table S1: **Composition of the mock community and reference of strains.**

| Expected | UNITE_s | | | | UNITE | | | | UNITE+INSD | | | |
|---|---|---|---|---|---|---|---|---|---|---|---|---|
| | Genus | Bootstrap | Species | Bootstrap | Genus | Bootstrap | Species | Bootstrap | Genus | Bootstrap | Species | Bootstrap |
| A. fischeri | Aspergillus | 100 | NA | 22 | Aspergillus | 100 | NA | 43 | Aspergillus | 100 | NA | 46 |
| A. fumigatus | Aspergillus | 100 | NA | 43 | Aspergillus | 98 | NA | 20 | Aspergillus | 100 | A. fumigatus | 68 |
| C. albicans | Candida | 99 | C. albicans | 89 | Candida | 100 | C. albicans | 100 | Candida | 100 | C. albicans | 70 |
| C. glabrata | Nakaseomyces | 100 | NA* | 100 | Nakaseomyces | 100 | NA* | 100 | Nakaseomyces | 94 | C. glabrata | 94 |
| C. metapsilosis | Candida | 100 | C. metapsilosis | 93 | Candida | 100 | C. metapsilosis | 100 | Candida | 100 | C. metapsilosis | 100 |
| C. parapsilosis | Candida | 100 | C. parapsilosis | 99 | Candida | 100 | C. parapsilosis | 55 | Candida | 100 | C. parapsilosis | 100 |
| C. parapsilosis | Candida | 100 | C. parapsilosis | 99 | Candida | 100 | C. parapsilosis | 58 | Candida | 100 | C. parapsilosis | 100 |
| M. carribica | Meyerozyma | 100 | M. guilliermondii | 96 | Meyerozyma | 100 | M. guilliermondii | 91 | Meyerozyma | 100 | M. carpophila | 90 |
| M. guillermondii | Meyerozyma | 100 | M. guilliermondii | 98 | Meyerozyma | 100 | M. guilliermondii | 100 | Meyerozyma | 100 | M. guilliermondii | 51 |
| M. sympodialis | Malassezia | 100 | M. sympodialis | 100 | Malassezia | 100 | M. sympodialis | 100 | Malassezia | 100 | M. sympodialis | 100 |
| S. cerevisiae | Saccharomyces | 100 | S. cerevisiae | 91 | Saccharomyces | 100 | S. cerevisiae | 95 | Saccharomyces | 100 | Saccharomyces species** | 100 |
| S. cerevisiae | Saccharomyces | 100 | S. cerevisiae | 100 | Saccharomyces | 100 | S. cerevisiae | 96 | Saccharomyces | 100 | Saccharomyces species** | 100 |

Table S2: Comparison of taxonomic annotation by three versions of the UNITE database and the *assignTaxonomy* function of dada2 by using a seed of 200. UNITE_s: UNITE database including global and 97% singletons; UNITE: UNITE database including singletons set as reference sequences; UNITE+INSD: full UNITE and INSD database. * Top hit is a sequence without Species-level annotation in UNITE. ** Top hit is a sequence with an undefined Species-level annotation in UNITE.

| Expected | e-value | Genus | Species |
|---|---|---|---|
| A. fischeri | 2.60e-166 | Aspergillus | A. oerlinghausensis |
| A. fumigatus | 2.60e-166 | Aspergillus | A. fumigatus |
| C. albicans | 1.08e-129 | Candida | C. africana |
| C. glabrata | 0 | Nakaseomyces | C. glabrata |
| C. metapsilosis | 4.09e-139 | Candida | C. metapsilosis |
| C. parapsilosis | 2.38e-136 | Candida | C. parapsilosis |
| C. parapsilosis | 5.13e-138 | Candida | C. parapsilosis |
| M. sympodialis | 5.42e-133 | Malassezia | M. sympodialis |
| M. caribbica | 1.92e-157 | Meyerozyma | M. caribbica |
| M. guillliermondii | 1.94e-152 | Meyerozyma | M. caribbica |
| S. cerevisiae | 0 | Saccharomyces | S. arboricola |
| | | Saccharomyces | S. bayanus |
| | | Saccharomyces | S. cariocanus |
| | | Saccharomyces | S. cerevisiae |
| | | Saccharomyces | S. eubayanus |
| | | Saccharomyces | S. kudriavzevii |
| | | Saccharomyces | S. mikatae |
| | | Saccharomyces | S. paradoxus |
| | | Saccharomyces | S. pastorianus |
| | | Saccharomyces | S. uvarum |
| S. cerevisiae | 0 | Saccharomyces | S. arboricola |
| | | Saccharomyces | S. bayanus |
| | | Saccharomyces | S. cariocanus |
| | | Saccharomyces | S. cerevisiae |
| | | Saccharomyces | S. eubayanus |
| | | Saccharomyces | S. kudriavzevii |
| | | Saccharomyces | S. mikatae |
| | | Saccharomyces | S. paradoxus |
| | | Saccharomyces | S. pastorianus |
| | | Saccharomyces | S. uvarum |

Table S3. Tied top hits with lowest e-value returned from a BLAST-based algorithm with the NCBI NT dataset as training set. By the nature of the database, each Species-level annotation has a unique representative sequence available in the NCBI NT database.

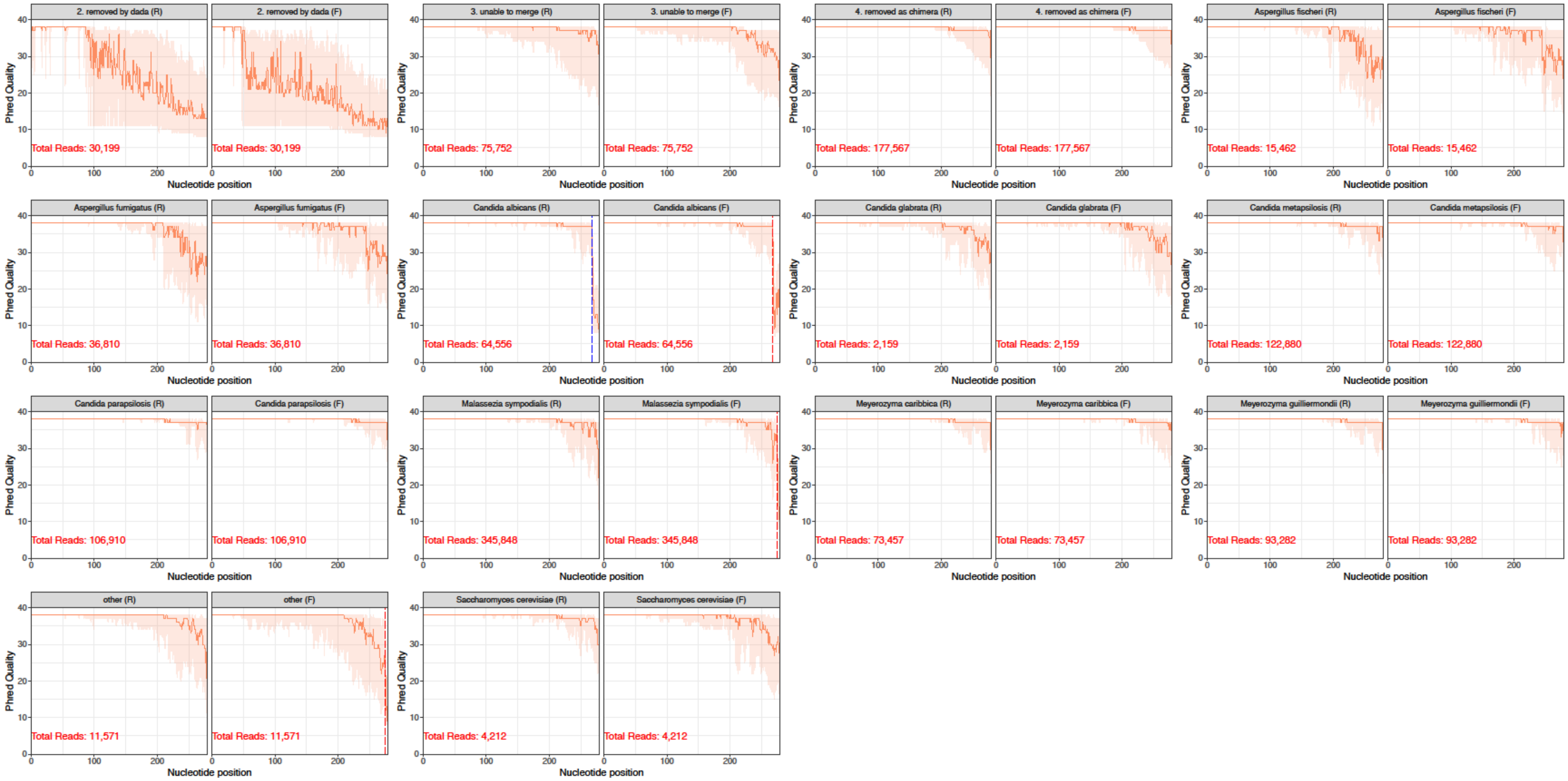


**Figure S1. Quality profile of raw reads with different subsequent fates in the DADA2 pipeline using maxEE=Inf, truncQ=0.** The orange line represents the median Phred Score at that position, while the shaded area represents the 25th to 75th percentiles. Vertical lines show the median length of reads after primer removal.

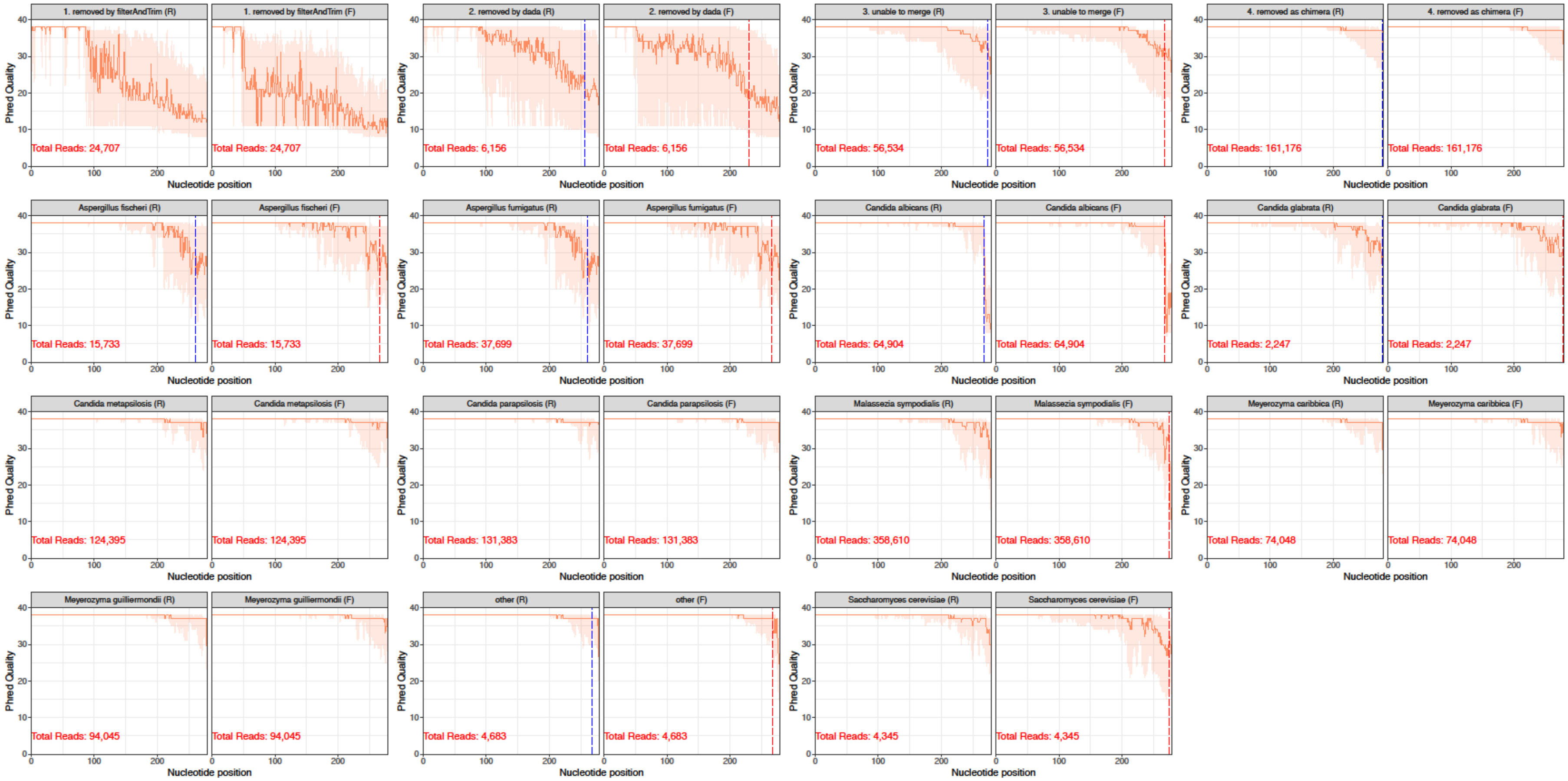


**Figure S2. Quality profile of raw reads with different subsequent fates in the DADA2 pipeline using maxEE=8, truncQ=8.** The orange line represents the median Phred Score at that position, while the shaded area represents the 25th to 75th percentiles. Vertical lines show the median length of reads after primer removal and quality truncation.
